# Corona exchange dynamics on carbon nanotubes by multiplexed fluorescence monitoring

**DOI:** 10.1101/761296

**Authors:** Rebecca L. Pinals, Darwin Yang, Alison Lui, Wendy Cao, Markita P. Landry

**Author notes:** Corresponding Author: Markita P. Landry. Author Contributions: These authors contributed equally (Rebecca L. Pinals, Darwin Yang).

## Abstract

Noncovalent adsorption of DNA on nanoparticles has led to their widespread implementation as gene delivery tools and optical probes. Yet, the behavior and stability of DNA-nanoparticle complexes once applied in biomolecule-rich, *in vivo* environments remains unpredictable, whereby biocompatibility testing usually occurs in serum. Here, we demonstrate time-resolved measurements of exchange dynamics between solution-phase and adsorbed corona-phase DNA and protein biomolecules on single-walled carbon nanotubes (SWCNTs). We capture real-time binding of fluorophore-labeled biomolecules, utilizing the SWCNT surface as a fluorescence quencher, and apply this corona exchange assay to study protein corona dynamics on ssDNA-SWCNT-based dopamine sensors. We study exchange of two blood proteins, albumin and fibrinogen, adsorbing to and competitively displacing (GT)_6_ vs. (GT)_15_ ssDNA from ssDNA-SWCNTs. We find that (GT)_15_ binds to SWCNTs with a higher affinity than (GT)_6_ and that fibrinogen interacts with ssDNA-SWCNTs more strongly than albumin. Albumin and fibrinogen cause a 52.2% and 78.2% attenuation of the dopamine nanosensor response, coinciding with 0.5% and 3.7% desorption of (GT)_6_, respectively. Concurrently, the total surface-adsorbed fibrinogen mass is 168% greater than that of albumin. Binding profiles are fit to a competitive surface exchange model which recapitulates the experimental observation that fibrinogen has a higher affinity for SWCNTs than albumin, with a fibrinogen on-rate constant 1.61-fold greater and an off-rate constant 0.563-fold smaller than that of albumin. Our methodology presents a generic route to assess real-time corona exchange on nanoparticles in solution phase, and more broadly motivates testing of nanoparticle-based technologies in blood plasma rather than the more ubiquitously-tested serum conditions.

## 1. INTRODUCTION

Adsorption of polymers on single-walled carbon nanotubes (SWCNTs) has enabled developments in molecular sensing,^1^ *in vivo* imaging,^2^ genetic cargo delivery,^3^ and chirality sorting.^4^ Noncovalent SWCNT functionalization offers a route that preserves the pristine atomic structure, thus retaining the intrinsic near-infrared (nIR) fluorescence of the SWCNTs for the aforementioned applications. However, noncovalent adsorption is an inherently dynamic process, where exchange occurs between molecules in the bulk solution and molecules on the surface, into what is known as the ‘corona phase’. In the case of polymers on SWCNTs, the nature, strength, and kinetics of both the polymer binding and unbinding processes are key contributors to the success of polymer-SWCNT based technologies.^5^ Understanding this exchange process is especially critical for intended uses of functionalized SWCNTs to probe biological environments. When a nanoparticle is injected into a biological system, the nanoparticle surface is spontaneously and rapidly coated with proteins to form the ‘protein corona’.^6^ In the case of noncovalent polymer-SWCNT complexes, we hypothesize that native biomolecules compete with the original polymer to occupy the nanoparticle surface. Binding of proteins and other biomolecules to the SWCNT can disrupt the intended functionality of the nanoparticle and lead to potentially adverse biocompatibility outcomes.^7,8^ This phenomenon of protein corona formation leads to challenges in translating *in vitro* sensing or biomolecule delivery platforms to *in vivo* application. Moreover, the generally accepted method of simulating *in vivo* biological conditions involves testing nanotechnology performance in blood serum.^2,9^ Yet, the absence of blood coagulation proteins from serum could yield a false outcome in assessing robustness of the nanotechnology and accordingly result in unpredicted failure when applied *in vivo*.

To clarify how nanoparticle-polymer conjugates behave in biologically-relevant environments, it is pivotal to understand the kinetics describing molecular exchange on nano-particle surfaces. Hence, we aim to gain a mechanistic understanding of how SWCNT-based neuromodulator sensors behave in protein-rich milieus. These sensors are based on noncovalent functionalization of (GT)_6_ single-stranded DNA (ssDNA) on SWCNTs, resulting in a complex that exhibits ultrasensitive ΔF/F_o_ = 2400% and 3500% fluorescence “turnon” responses in the presence of neuromodulators dopamine and norepinephrine, respectively.^10–12^ However, the drastic enhancement of SWCNT fluorescence experienced upon *in vitro* exposure to dopamine is attenuated to ΔF/F_o_ ≈ 20% once the sensors are applied in brain tissue,^12^ presumably due to protein adsorption and/or disruption of the ssDNA corona phase originally on the SWCNT surface.

Current methods to measure dynamic, noncovalent exchange on nanoparticles exist but are limited in scope. Most research on protein-surface interactions involves characterizing macroscopic surfaces using a series of well-developed techniques that broadly entail an input signal modulated by changing adsorbate mass on the surface as a function of time, including total internal reflection fluorescence microscopy, surface plasmon resonance, biolayer interferometry, and quartz-crystal microbalance with dissipation monitoring. To apply these surface techniques to nanoparticles, the nanoparticles must be surface-immobilized, thus introducing unrealistic topographical constraints that affect ligand exchange kinetics, lead to mass transport limitations,^13^ do not reproduce solution-phase nanosensor responses,^14^ and cause nonselective protein adsorption to any surface left exposed during the sparse SWCNT immobilization process.^14^

An alternative method that permits the study of SWCNTs in solution takes advantage of SWCNT sensitivity to their local dielectric environment^15–17^ by monitoring SWCNT fluorescence intensity changes and solvatochromic shifts upon corona exchange.^18,19^ This technique is applied to study polymer-surfactant exchange kinetics,^20–23^ whereby SWCNTs suspended with surfactant exhibit higher quantum yield and optical transition energy (i.e. blue-shifted spectra) compared to SWCNTs suspended with most biomolecules such as protein or ssDNA. Previous work has successfully applied measurable differences in SWCNT fluorescence spectra to study relative changes in corona surface composition.^24,25^ However, this approach cannot distinguish the exchange of two biomolecules (here, ssDNA to protein), nor can it distinguish between molecular rearrangement vs. molecular desorption from the SWCNT surface. Despite the advantage of undertaking corona exchange studies in the solution phase with this approach, its low sensitivity, non-quantitative nature, and inability to distinguish between adsorbed biomolecules nullifies its potential for monitoring ssDNA-protein exchange.

In this work, we present an assay that overcomes the limitations of previous characterization methods to study corona exchange dynamics between solution-phase and corona-phase biopolymers on SWCNTs, specifically applied to ssDNA and protein. This assay exploits the quenching of fluorophores when in close proximity to the SWCNT surface to monitor ligand binding and unbinding events.^26^ While prior literature has similarly harnessed fluorophore quenching by SWCNTs to study the ssDNA-to-SWCNT binding process,^8,18,27^ far less is known regarding how pre-adsorbed ssDNA and biologically native proteins exchange on the SWCNTs. To our knowledge, this method is unique in enabling real-time monitoring of SWCNT surface exchange between ssDNA and proteins, tracing the fate of all biomolecules involved in the binding exchange. We conduct multiplexed fluorescence tracking of polymer adsorption and desorption events to/from the SWCNT surface. As a case study for this assay, we focus on comparing the sorption behavior of two specific blood proteins, human serum albumin and fibrinogen, chosen because: (i) both are highly abundant in plasma, with albumin as ~55% (w/v) of blood plasma, or 35-50 mg/mL^28^ and fibrinogen as ~4% (w/v) of blood plasma, or 1.5-4.5 mg/mL^29^, (ii) albumin is present in both blood plasma and serum, whereas fibrinogen is a key coagulation protein present in plasma but depleted from serum, and (iii) albumin and fibrinogen are known to be interfacially active proteins prone to surface-adsorption and are implicated in the formation of many other nanoparticle coronas.^30–32^ Binding profiles from the experimental assay in conjunction with a competitive-exchange model are used to extract kinetic parameters for each adsorbent species. Although this study specifically examines competitive adsorption of individual plasma proteins, albumin and fibrinogen, onto (GT)_6_- and (GT)_15_-SWCNTs, the assay is general to any molecules that can be fluorescently labeled and to any nanomaterial surface to which these species may adsorb and display quenched fluorescence. Binding is also validated against the orthogonal and more ubiquitously-used platform monitoring solvatochromic shifting of the nIR SWCNT spectrum as a proxy for SWCNT corona coverage.^24,25^ The work presented herein develops an understanding of the fundamental corona exchange mechanism, contextualizes the nature of the ligand exchange process vs. SWCNT solvatochromic shifting, and provides guidance for testing the performance of SWCNT-based systems in biologically relevant, protein-rich conditions.

## 2. RESULTS AND DISCUSSION

### 2.1 Proteins attenuate dopamine sensor response

Noncovalent modification of single-walled carbon nanotubes (SWCNTs) with single-stranded (GT)_6_ DNA imparts nIR fluorescence responsivity to the small molecule neurotransmitter, dopamine.^10,11,33^ Addition of 200 μM dopamine to 5 μg/mL solution-phase (GT)_6_-SWCNTs in phosphate buffered saline (PBS) yields an 11.5-fold increase in nanosensor fluorescence at the 1200 nm SWCNT emission peak (Fig. 1a; see **4.4 Near-infrared fluorescence measurements** methods section). Nanosensor response was diminished in the presence of 40 μg/mL human serum albumin (HSA, Fig. 1b) and 40 μg/mL fibrinogen (FBG, Fig. 1c), proteins abundant in intravenous environments. Incubation of 40 μg/mL HSA or FBG with 5 μg/mL (GT)_6_-SWCNTs reduced fluorescence dopamine response by 52.2% or 78.2% after 40 minutes (Fig. 1d), respectively. Protein-induced attenuation of SWCNTs. HSA did not cause any wavelength shifting of the (GT)_6_-SWCNT emission, while FBG exposure led to a redshift of 2.6 ± 0.6 nm (mean ± standard deviation of N=3 sample replicates). Although changes in both the nIR fluorescence intensity and emission wavelength could indicate protein binding, monitoring the SWCNT fluorescence alone does not provide sufficient information to correlate these phenomena.

**Figure 1.**
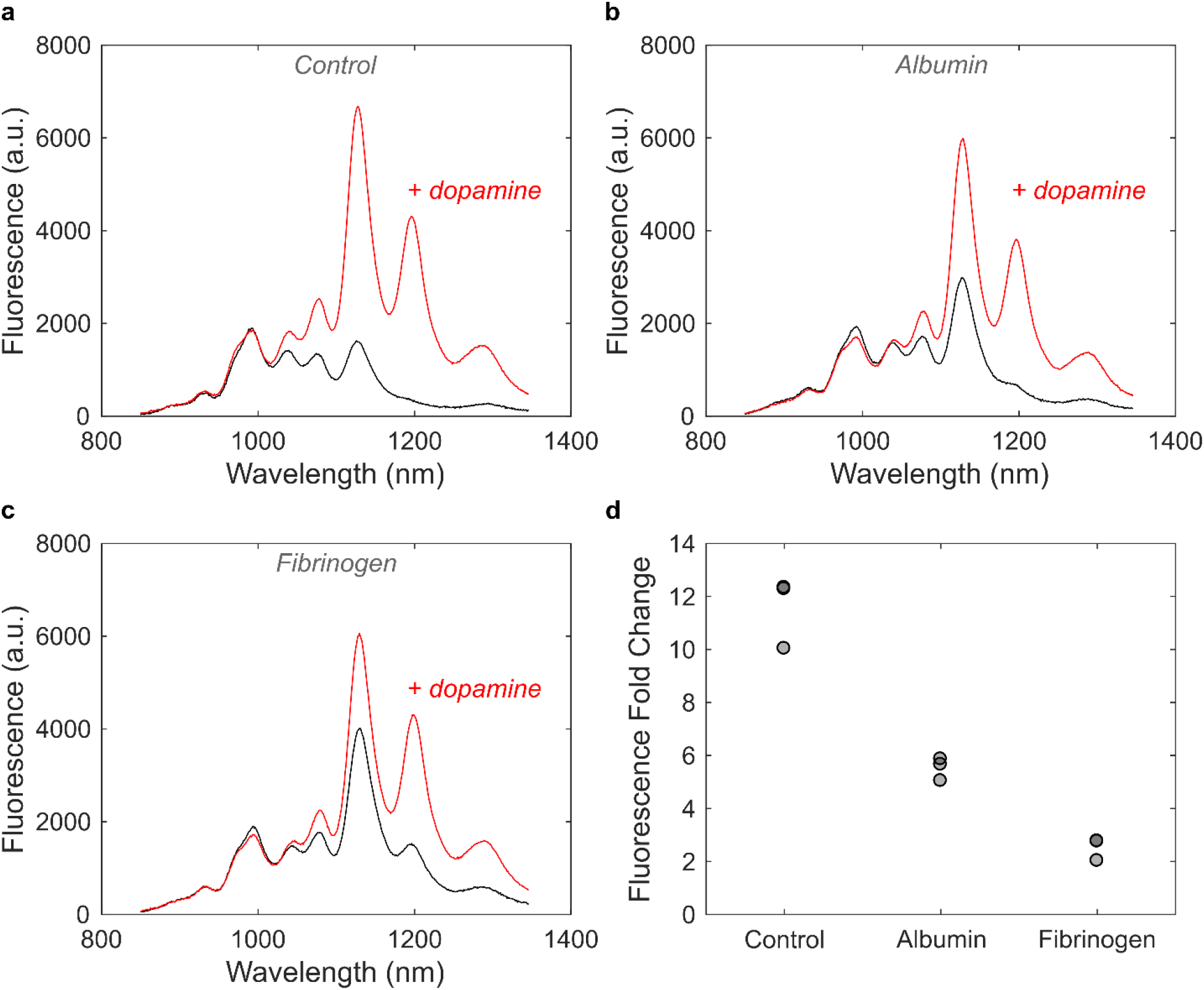
Protein adsorption attenuates (GT)_6_-SWCNT sensor response to dopamine. **(a)** Near-infrared (nIR) spectra of 5 μg/mL (GT)_6_-SWCNTs before (black) and after (red) addition of 200 μM dopamine. **(b-c)** nIR spectra of 5 μg/mL (GT)_6_-SWCNTs incubated with 40 μg/mL **(b)** albumin or **(c)** fibrinogen for 40 minutes before (black) and after (red) addition of 200 μM dopamine. **(d)** Change in (GT)_6_-SWCNT fluorescence intensity at 1200 nm peak following 40 minutes incubation in PBS or protein solution at 40 μg/mL, then addition of 200 μM dopamine (N = 3). Nanosensor excitation was with 721 nm light.

We first implemented the solvatochromic shift assay to study surfactant-induced fluorescence changes of 5 μg/mL (GT)_6_-SWCNTs incubated with either 40 μg/mL HSA or FBG for 40 minutes. Displacement of the biopolymer corona phase with surfactant, here 0.5% (w/v) sodium dodecylbenzenesulfonate (SDBS), causes a change in local dielectric environment that in turn leads to a blue shift in SWCNT emission wavelength and an increase in SWCNT fluorescence emission intensity. The magnitude of these observed effects is thought to provide insight on the original SWCNT-corona stability. Interestingly, FBG incubated with (GT)_6_-SWCNTs resulted in both the largest magnitude wavelength shift and largest fold change in fluorescence intensity upon addition of SDBS (Fig. S1). In contrast, HSA incubated with (GT)_6_-SWCNTs did not show a significantly different wavelength shift or intensity fold change compared to the control, (GT)_6_-SWCNTs incubated with only PBS. These results suggest albumin and fibrinogen proteins may have different binding propensities and kinetics to the SWCNT surface. However, this test fails to decouple the interactions between SWCNTs with ssDNA, protein, and then surfactant. To further study the differential attenuation of sensor response by HSA and FBG, and more thoroughly understand the exchange dynamics occurring on the SWCNT surface, we developed a method for studying SWCNT corona composition by multiplexed fluorescence monitoring.

### 2.2 Multiplexed fluorescence tracking enables realtime monitoring of ligand exchange dynamics

Our assay leverages fluorophore quenching induced by proximity to the SWCNT surface to measure surface exchange dynamics. Proteins under study were labeled with a FAM fluorophore (ex/em = 494/520 nm) using NHS ester conjugation to primary amine groups (see **4.2 Fluorophore-labeling of proteins** methods section). Single-stranded DNA (ssDNA) were procured with a 3’ terminally-labeled Cy5 fluorophore (ex/em = 648/668 nm), enabling spectrally resolved multiplexed tracking of protein and ssDNA. The ssDNA-Cy5 is initially quenched on the SWCNT surface, increasing in fluorescence upon desorption from the SWCNT. Conversely, the FAM-labeled protein exhibits high fluorescence when added in bulk solution, quenching upon adsorption to the SWCNT surface. In this manner, FAM-labeled protein can be injected into ssDNA-Cy5-SWCNTs in a well-plate format and fluorescence changes resulting from biomolecule exchange can be read by a fluorescence plate reader (Fig. 2a). We first employed this method to compare the desorption rates of (GT)_6_-Cy5 and (GT)_15_-Cy5 from SWCNTs upon addition of FAM-labeled HSA and FBG. Both proteins promoted dequenching of Cy5, as compared to the addition of PBS control (Fig. 2b-c). Dequenching was due to complete desorption of ssDNA rather than partial desorption of the 3’ end, as verified by confirming that the binding profiles of 3’-vs. internally Cy5-labeled ssDNA are similar (Fig. S2). Fibrinogen generated a 3.09 ± 0.07 Cy5 fluorescence fold increase for (GT)_6_-Cy5-SWCNTs vs. a 1.52 ± 0.04 Cy5 fluorescence fold increase for (GT)_15_-Cy5-SWCNTs. This result suggests (GT)_15_ is less readily displaced from the SWCNT surface compared to the shorter (GT)_6_ construct, a result consistent with the literature.^34–36^

**Figure 2.**
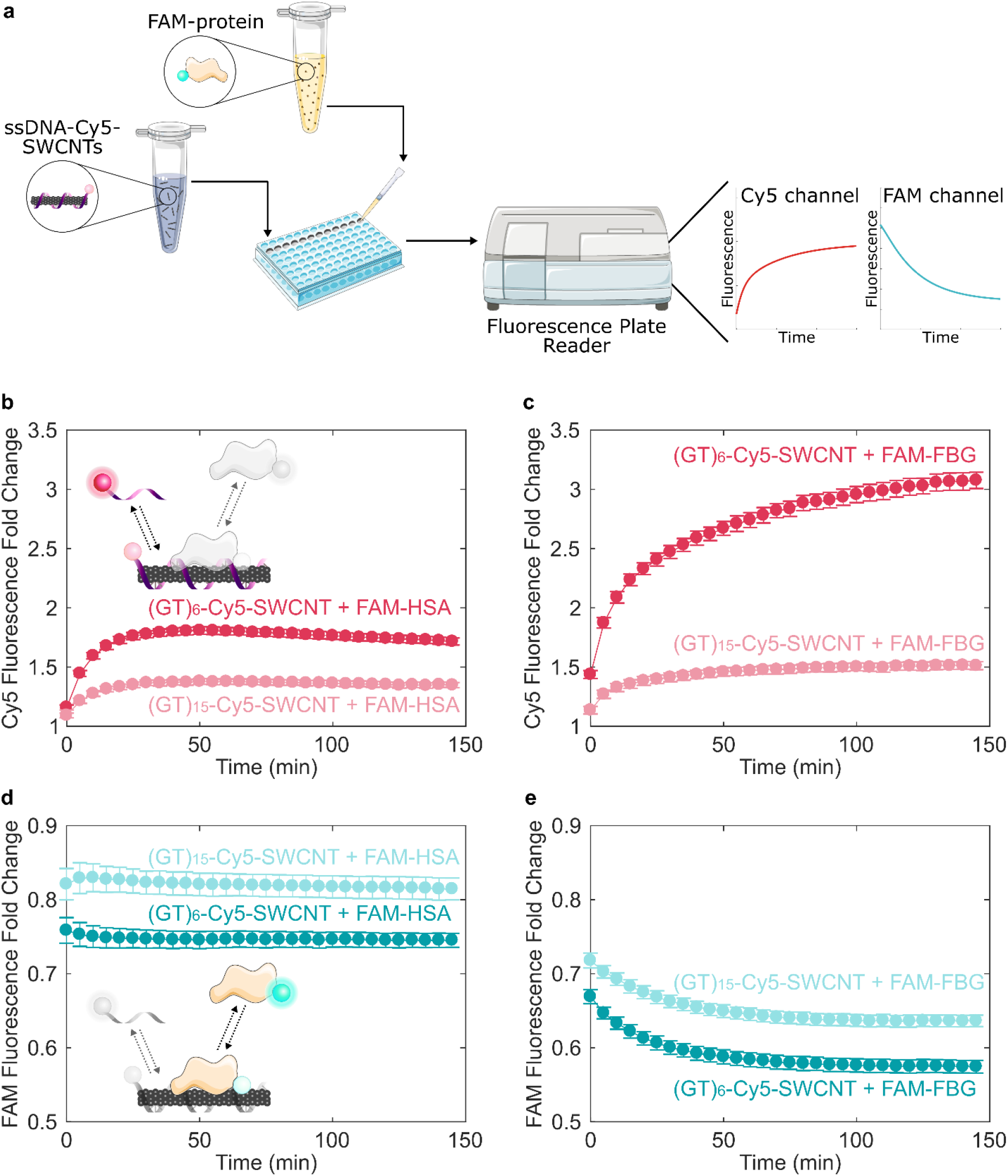
Tracking exchange of fluorophore-labeled ssDNA and protein on SWCNT surfaces demonstrates protein adsorption selectivity and ssDNA length effect. **(a)** Corona exchange assay workflow. ssDNA-Cy5-SWCNT solution was added to a well-plate, FAM-protein solution was injected, and the ad-/de-sorption processes were monitored in separate color channels of a fluorescence plate reader (see **4.3 Visible fluorescence measurements** methods section). Increase in ssDNA-Cy5 fluorescence induced by addition of 40 μg/mL **(b)** FAM-labeled albumin (FAM-HSA) or **(c)** FAM-labeled fibrinogen (FAM-FBG) to 5 μg/mL ssDNA-Cy5-SWCNT suspended with ssDNA, (GT)_6_ or (GT)_15_. Decrease in fluorescence of **(d)** FAM-HSA and **(e)** FAM-FBG after addition of protein to (GT)_6_- or (GT)_15_-SWCNT. Error bars represent standard error between experimental replicates (N = 3).

In the same experiment, protein adsorption onto ssDNA-Cy5-SWCNTs was concurrently tracked via fluorescence quenching of the protein-conjugated FAM. Presence of residual FAM fluorophore in the FAM-protein solution was accounted for by quantifying free FAM and subtracting the minimal change in fluorescence due to free FAM-to-SWCNT interaction (Fig. S3, S4, Table S1). Furthermore, the effect of FAM fluorophore labeling on the protein-exchange dynamics was minimal (Fig. S5), in agreement with previous investigations demonstrating that fluorescein-labeling of proteins does not perturb protein adsorption or function, and additionally that fluorescein signals are proportional to the interfacial mass of the tagged species. ^37–40^ By tracking the fluorescence modulation resulting from FAM-protein interactions with ssDNA-Cy5-SWCNTs, we found that FAM-FBG exhibited a comparatively larger degree of quenching than FAM-HSA for both ssDNA-SWCNT suspensions (Fig. 2d-e): upon addition of 40 μg/mL FAM-FBG to 5 μg/mL (GT)_6_-SWCNTs (final concentrations), FBG induced a 42.5 ± 0.9% decrease in FAM fluorescence vs. a 25.5 ± 0.9% HSA-induced decrease in FAM fluorescence. These results consistently suggest two interaction mechanisms of ssDNA and protein with SWCNTs: (i) (GT)_15_ ssDNA binds to SWCNTs with a higher affinity than (GT)_6_ ssDNA, thus reducing protein adsorption, and (ii) FBG interacts with ssDNA-SWCNTs more strongly than HSA. The former result agrees with prior work confirming that the rate of ssDNA desorption from SWCNTs decreases with increasing oligo length,^41^ also valid in the presence of competing biomolecules. ^8^ As such, our data suggest that FBG protein adsorption leads to more significant ssDNA desorption from SWCNTs, whereas HSA adsorbs less strongly and accordingly causes less ssDNA desorption from SWCNTs. Interestingly, protein adsorption occurs faster than ssDNA desorption. These experimental results motivate kinetic modeling of ssDNA and protein exchange on SWCNT surfaces.

### 2.3 Kinetic modeling of ssDNA/protein competitive binding on SWCNT surface

To quantitatively probe differences in protein affinities for ssDNA-SWCNTs, we fit Cy5 and FAM fluorescence data to a competitive adsorption model and extracted kinetic parameters for ssDNA and proteins. Multiplexed fluorescence tracking was repeated with 5 μg/mL (GT)_6_-Cy5-SWCNTs and concentrations of FAM-HSA and FAM-FBG ranging from 5 to 160 μg/mL. Fluorescence values were converted to mass concentration using standard curves for ssDNA-Cy5 and both FAM-conjugated proteins (Fig. S6). A model was developed for the competitive exchange between ssDNA and protein on the SWCNT surface (Equations 1 and 2). In the model, unbound ssDNA (D) and protein (P) adsorb and desorb reversibly to SWCNT surface sites (*):

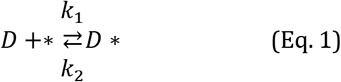

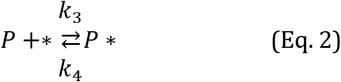

Total concentration of SWCNT surface sites ([*]_τ_) was fixed, given by a site balance (Equation 3), where *, D*, and P* refer to vacant sites, sites occupied by bound ssDNA, and sites occupied by bound protein, respectively:

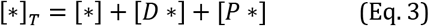

Bound ssDNA and bound protein concentrations were calculated by species conservation, where total ssDNA was determined by desorption experiments (see **4.1 Preparation of ssDNA-SWCNT complexes** methods section), total protein was the injected protein quantity, and total vacant sites ([*]_τ_) was a fit parameter. Rate constants k_1_, k_2_ for ssDNA binding/unbinding, k_3_, k_4_ for protein binding/unbinding, and the total concentration of binding sites [*]_T_ were computed using a least squares curve fit of Equations S1 and S2 to experimental data (see **4.5 Kinetic model** methods section).

Experimental data of FAM-HSA or FAM-FBG added to (GT)_6_-Cy5-SWCNTs was fit to this model for each concentration tested (Fig. 3). All mean relative errors comparing fits to experimental data were < 5% (Table S2). The model recapitulates the experimental observation that FBG has a higher affinity for SWCNTs (Fig. 3d) than HSA (Fig. 3b), where average k_3, FBG_ = 1.43 × 10^−5^ > k_3, HSA_ = 8.88 × 10^−6^ mL μg ^−1^ s^−1^ (Table 1, with full fit parameter results in Table S2). At the same initial FAM-protein concentration of 40 μg/mL added to 5 μg/mL (GT)_6_-Cy5-SWCNTs, FBG adsorbed to a higher fraction of bound protein (0.756) than HSA (0.284) after 1 hour. Addition of FAM-FBG into solution with (GT)_6_-Cy5-SWCNTs led to ssDNA desorption for all tested concentrations of injected FBG (Fig. 3c) but only led to measurable ssDNA desorption for concentrations ≥ 40 μg/mL of injected HSA (Fig. 3a). Adsorption of ssDNA was observed upon addition of PBS or low concentrations of FAM-HSA (≤ 20 μg/mL) to (GT)_6_-Cy5-SWCNTs, indicating an initial excess of unbound ssDNA in bulk solution. Interestingly, for all concentrations tested, protein adsorption proceeded significantly faster than ssDNA desorption dynamics, indicating that protein adsorption precedes ssDNA desorption and suggesting that the two phenomena may be decoupled in time. This difference in exchange timescales may be due to the large concentration of total SWCNT surface binding sites (with average fit values of [*]_T,FBG_ = 572 μg mL^−1^ and [*]_T,HAS_ = 472 μg mL^−1^) relative to the total ssDNA and protein concentrations. We hypothesize a low initial ssDNA surface coverage, or large accessible SWCNT surface area, is a likely reason for the difference in exchange timescales. Furthermore, in the case of FBG with (GT)_6_-Cy5-SWCNTs, while amount of adsorbed FBG reaches an apparent steady-state value within ~5 minutes (Fig. 3d), ssDNA continues to gradually desorb over time at a rate seemingly independent of injected protein concentration (Fig. 3c). Continued ssDNA desorption may be caused by a surface rearrangement process in the adsorbed FBG layer,^42^ where protein spreading could be responsible for this observed ssDNA displacement over longer timescales.^13,38^ Hydrophobic interactions are posited to be the driving force for protein spreading on the SWCNT surface^39^ and consequently interfacial denaturation is presumed to be the dominant relaxation process, in addition to a smaller contribution from molecular reorientations.^37^

**Table 1.**
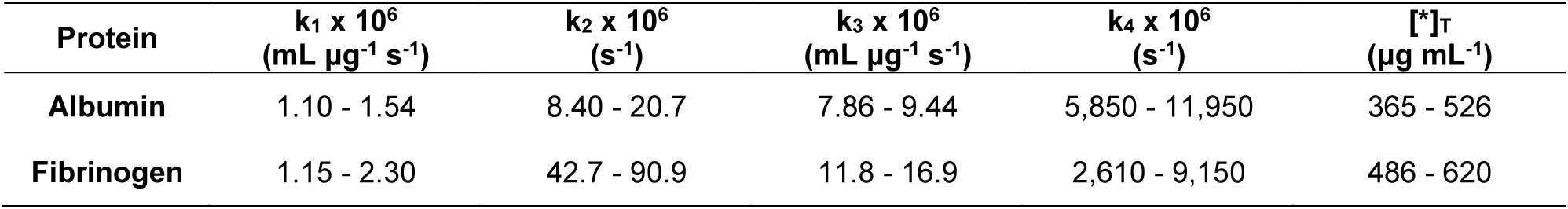
Range of kinetic model fit parameters.

**Figure 3.**
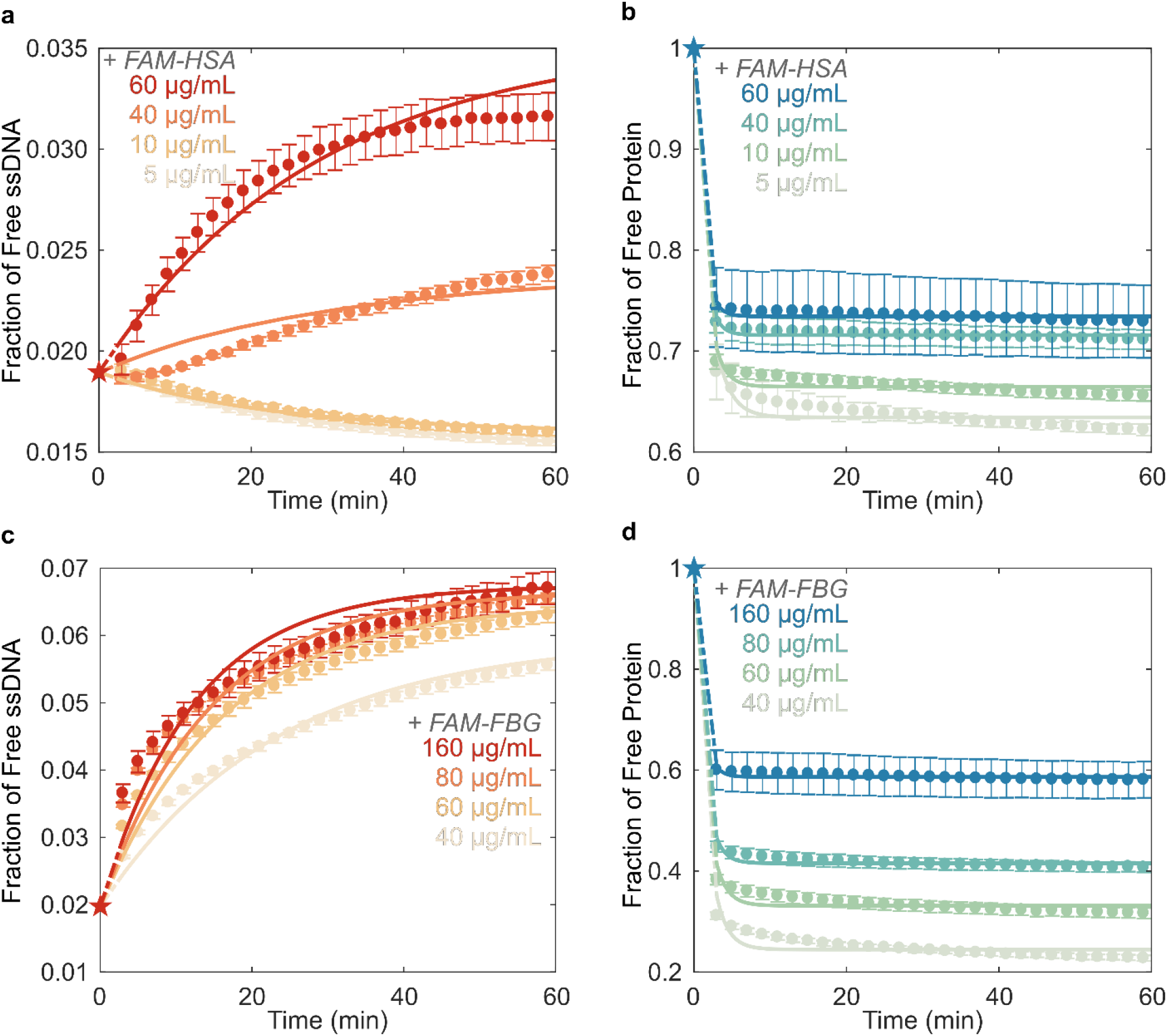
Kinetic model of competitive exchange between ssDNA and protein on SWCNTs fit to fluorescence data to extract rate constants. Fraction of **(a)** (GT)_6_-Cy5 ssDNA and **(b)** FAM-labeled albumin (FAM-HSA) protein free in solution for varying concentrations of FAM-HSA injected into 5 μg/mL (GT)_6_-Cy5-SWCNT solution. Fraction of **(c)** (GT)_6_-Cy5 ssDNA and **(d)** FAM-labeled fibrinogen (FAM-FBG) protein free in solution for varying concentrations of FAM-FBG injected into (GT)_6_-Cy5-SWCNT solution. Star data points represent initial conditions used for solving model differential equations. Error bars represent standard error between experimental replicates (N = 3).

From the kinetic model fitting, the mass of protein adsorbed on the SWCNTs was consistently higher for FBG as compared to HSA for the same initial concentration of 40 μg/mL FAM-protein with 5 μg/mL (GT)_6_-Cy5-SWCNTs (see **4.5 Kinetic model** methods section). Previous studies of differential protein adsorption to hydrophobic surfaces has demonstrated that fibrinogen forms well-packed layers, whereas more weakly adsorbed albumin forms less tightly packed, more mobile adsorption layers.^39^ Accordingly, we hypothesize that the seemingly higher protein surface coverage on the SWCNT points to the more tightly packed, if not multilayer formation, of FBG on the SWCNT surface.

Finally, we note that the dependence of rate constants k_2_ and k_4_ on protein concentration indicates that the model is not fully descriptive of the system (Table S2). Some potential shortcomings include that the proposed elementary steps only approximate the true exchange mechanism, or that there are nonidealities present in the protein and/or ssDNA sorption behavior. Nevertheless, the kinetic model captures the key binding profiles and provides a relative comparison between biomolecules binding to the SWCNT surface.

## 3. CONCLUSIONS

Protein adsorption to nanoparticle surfaces is a major hindrance to the successful application of nanotechnologies *in vivo*. We have shown that incubation of two high-abundance blood plasma proteins, human serum albumin and fibrinogen, with ssDNA-SWCNT dopamine sensors causes significantly different degrees of sensor response attenuation. Developing an understanding of protein-sensor interactions is vital in circumventing this issue and establishing better practices for testing nanotechnologies for *in vivo* use. Previously established techniques to evaluate these effects implement surface-immobilized nanoparticles or exploit the intrinsic nIR fluorescence changes of SWCNTs. Yet, these methods do not track the fate of adsorbates and cannot quantify the fraction of free biomolecules in real-time, thus precluding quantitative and temporally-resolved studies of the SWCNT protein corona composition. Though the SDBS-induced solvatochromic shift assay successfully identifies FBG as a protein of interest, this assay provides no mechanistic information on FBG binding, nor can it distinguish between HSA and PBS control responses.

We have addressed these limitations in developing a method to quantitatively probe the kinetics of SWCNT corona exchange between ssDNA and protein adsorbates by monitoring fluorescence quenching of conjugated fluorophores in close proximity to SWCNT surfaces. Concentration curves were fit to a competitive adsorption model to extract kinetic parameters. Our method reveals that reduction of dopamine sensor performance correlates with quantity of adsorbed protein, where fibrinogen adsorbs to ssDNA-SWCNTs 168% more than albumin at the same concentration, and consequently leads to 26% more sensor attenuation. We demonstrate significantly greater SWCNT binding affinities for longer repeating ssDNA sequences, and for fibrinogen over albumin. These results bear significance in that albumin is the highest abundance blood protein and is therefore commonly regarded as an important component of the SWCNT corona. However, our results show that lower abundance proteins with higher SWCNT affinities may disproportionately contribute to the SWCNT corona, as has been previously suggested in orthogonal protein corona-nanoparticle studies.^43,44^ Preliminary findings from blood plasma and serum samples normalized to 40 μg/mL total protein concentration show that plasma and serum both cause significant attenuation of dopamine response in (GT)_6_-Cy5-SWCNTs, with 81.0 ± 0.9% and 80.7 ± 1.4% reduction in response, respectively (Fig S7a). However, plasma—which contains fibrinogen—caused a higher degree of ssDNA desorption, with plasma inducing a 1.64 ± 0.01 fold increase in (GT)_6_-Cy5 fluorescence vs. a 1.39 ± 0.03 fold increase by serum (Fig S7b). Our results motivate the necessity to test SWCNT-based and other nanobiotechnologies in more representative bio-environments, i.e. blood plasma rather than serum.

Furthermore, the method presented herein enables the study of protein corona formation dynamics, with standard laboratory equipment, under varying solution conditions (e.g. ionic strength and pH) with multiple biomolecular entities. This in turn will facilitate more thorough deconvolution of factors driving protein corona formation and accordingly inform design principles for nanotechnologies resistant to protein corona-based biofouling and performance attenuation. Careful selection of fluorophores may enable further multiplexing, allowing tracking of more than two distinct molecular species simultaneously. Rationally designed labeling methodologies such as FRET may also enable the study of more complex interactions such as protein denaturation on the SWCNT surface. In summary, the corona exchange assay we have developed will serve to enhance our still deficient understanding of how noncovalently bound polymers exchange on nanoparticle surfaces and, accordingly, enable the design and testing of nanobiotechnologies towards effective implementation *in vivo*.

## 4. MATERIALS AND METHODS

### 4.1 Preparation of ssDNA-SWCNT complexes

Single-stranded DNA with single-walled carbon nanotube (ssDNA-SWCNT) suspensions were prepared by combining 0.2 mg of mixed-chirality SWCNTs (small diameter HiPco™ SWCNTs, NanoIntegris) and 20 μM of ssDNA (custom ssDNA oligos with standard desalting, Integrated DNA Technologies, Inc.) in 1 mL of 0.01 M phosphate-buffered saline (PBS). Solutions were probe-tip sonicated for 10 minutes in an ice bath (3 mm probe tip set at 50% amplitude, 5-6 W, Cole-Parmer Ultrasonic Processor). Samples were centrifuged to pellet insoluble SWCNT bundles and contaminants (16,100 cfg for 30 minutes). Supernatant containing the product was collected. ssDNA-SWCNT solutions were stored at 4°C until use. ssDNA-SWCNT concentration was determined via sample absorbance at 910 nm and the corresponding extinction coefficient ε_910nm_ = 0.02554 mL μg^-1^ cm^-1^.^45^ ssDNA-SWCNTs were diluted to a working concentration of 10 μg/mL in 0.1 M PBS.

Cyanine 5 (Cy5) was chosen as the ssDNA fluorophore label, with excitation maximum at 648 nm and emission maximum at 668 nm. The same suspension protocol was employed for preparation of fluorophore-labeled ssDNA-SWCNT complexes, using ssDNA-Cy5 (3’ or internally-labeled Cy5-labeled custom ssDNA oligos with HPLC purification, Integrated DNA Technologies, Inc.) in place of unlabeled ssDNA. Internally labeled ssDNA-Cy5 includes Cy5 conjugated to the thymine at nucleotide position 6 (GTGTGT/iCy5/GTGTGT).

Total ssDNA adsorbed to SWCNTs was determined by a heat/surfactant elution process. This molar ratio of ssDNA:SWCNT was required to calculate the fraction of free vs. bound ssDNA throughout the exchange process. Optimized elution conditions were achieved with salt and surfactant in the combination of 0.1 M PBS/0.1% (m/v) sodium dodecylbenzenesulfonate (SDBS), in agreement with prior literature demonstrating that SDBS disperses SWCNTs most effectively^18,46^ and without chirality bias.^47^ Freshly prepared ssDNA-Cy5-SWCNTs were diluted to a final concentration of 5 μg/mL in elution buffer, with a final volume of 150 μL in a PCR tube. Samples were heated at 95℃ for 1 hour in a PCR thermocycler, transferred to a clean test-tube, and centrifuged (16,100 cfg for 30 minutes) to pellet insoluble SWCNT bundles. 120 μL of supernatant containing the eluted ssDNA-Cy5 was collected. Fluorescence in the Cy5 channel was measured (see **4.3 Visible fluorescence measurements** methods section) and compared to a standard curve of known ssDNA-Cy5 concentrations (ranging 0.01 – 1 μM) to correlate the Cy5 fluorescence measurement to ssDNA eluted concentration. This resulted in a mole ratio of 364.20 ± 2.08 (GT)_6_:SWCNT and 139.96 ± 6.67 (GT)_15_:SWCNT (both N = 8), in relative agreement with previous literature for (6,5) single chirality SWCNTs.^48^

### 4.2 Fluorophore-labeling of proteins

N-Hydroxysuccinimide (NHS) ester chemistry was used to label proteins via conjugation to primary amine groups. Fluorescein (FAM) was chosen as the protein fluorophore label, with excitation maximum at 494 nm and emission maximum at 520 nm (FAM NHS ester 6-isomer, Lumiprobe). Lyophilized proteins were purchased: human serum albumin (HSA; from human plasma, ≤0.02% Fatty acids, Lot #SLBZ2785, Millipore Sigma) and fibrinogen (FBG; from human plasma, 20 mM sodium citrate-HCl pH 7.4, Lot #3169957, Millipore Sigma). FAM-proteins were prepared with 10 mg of protein reconstituted in 900 μL of 0.1 M PBS and 8-fold molar excess of FAM NHS ester solubilized in 100 μL DMSO. Solutions were combined, covered in foil, and incubated on a test-tube rotator for 4 hours. FAM-protein conjugates were twice purified to remove free FAM (Zeba™ 2 mL spin desalting columns with 40 kDa MWCO, Thermo Fisher Scientific) by washing with 0.1 M PBS three times (1,000 cfg for 2 minutes), centrifuging with sample (1,000 cfg for 3 minutes), and retaining sample in flow-through solution (repeating all steps twice with a new spin column). Protein concentration and degree of labeling (DOL) were determined by measuring the absorbance of the FAM-protein conjugate at the protein absorbance maximum, 280 nm (A_280_), and the fluorophore emission maximum, 494 nm (A_494_). Protein absorbance was corrected for the contribution of the fluorophore to A280 by subtracting A_494_ weighted by the correction factor (CF), an empirical constant of 0.17 for free FAM (from manufacturer). Protein and FAM concentrations were determined by the Beer-Lambert Law using either A_280_ for protein or A_494_ for FAM, with the corresponding extinction coefficients of ε_280nm,HSA_ = 43,824 (M cm)^−1^,^49^ ε_280nm,FBG_ = 513,400 (M cm)^−1^,^50^ and ε_494nm,FAM_ = 75,000 (M cm)^−1^ (from manufacturer). DOL was then calculated as the ratio of FAM to protein molar concentrations, yielding DOL_FAM-HSA_ = 2.773 and DOL_FAM-FBG_ = 0.608.

Free FAM NHS ester remaining after purification was quantified by polyacrylamide gel electrophoresis (PAGE) run according to the Laemmli protocol^51^ (adapted in BioRad Mini-PROTEAN Tetra Cell manual). Briefly, purified FAM-protein conjugates were added to sodium dodecyl sulfate (SDS) reducing buffer (2% SDS, 5% β-mercaptoethanol, 0.066 M Tris-HCl) in a 1:2 ratio of sample to buffer. Samples were diluted such that 100 ng of FAM-HSA, 100 ng of FAM NHS ester, or 30 ng of FAM-FBG (due to lower labeling reaction yield) per 20 μL volume was applied per well. PAGE separation was carried out in 1 mm vertical mini gel format (Bio-Rad Mini-PROTEAN Tetra Cell) with a discontinuous buffer system under denaturing conditions. Gel composition was 12% acrylamide (total monomer), 0.375 M Tris-HCl, 0.1% SDS, 0.05% APS, 0.05% TEMED for the resolving gel and 12% acrylamide (total monomer), 0.125 M Tris-HCl, 0.1% SDS, 0.05% APS, 0.1% TEMED for the stacking gel. Electrode buffer was 25 mM Tris, 192 mM glycine, and 3.5 mM SDS (pH 8.3). Separation was run with 200 V for 35 minutes, gels were extracted, and the FAM label was visualized with a gel imager (Typhoon FLA 9500, 473 nm laser, General Electric) (Fig. S3). The FAM-protein conjugate is the higher band (approximately 66 kDa for FAM-HSA, 52-95 kDa for FAM-FBG) and the free, lighter molecular weight FAM NHS ester is the lower band (approximately 0.475 kDa). FAM fluorescence intensity was quantified with ImageJ (Table S1).

### 4.3 Visible fluorescence measurements

Equal volumes of (GT)_6_-Cy5-SWCNT and FAM-tagged protein at 2X working concentration were added to a 96-well PCR plate (Bio-Rad) to a total volume of 50 μL. The plate was sealed using an optically transparent adhesive seal (Bio-Rad) and briefly spun down on a benchtop centrifuge. Fluorescence time series readings were taken using a Bio-Rad CFX96 Real Time qPCR System, scanning all manufacturer set color channels (FAM, HEX, Texas Red, Cy5, Quasar 705) every 30 s at 22.5 °C (lid heating off). Fluorescence time series were analyzed without default background correction. Note that concentration ranges of FAM-HSA (5-60 μg/mL) and FAM-FBG (40-160 μg/mL) were chosen to be in the linear fluorescence regime of the qPCR.

### 4.4 Near-infrared fluorescence measurements

Fluorescence spectra were collected with an inverted Zeiss microscope (20x objective, Axio Observer.D1) containing a custom filter cube set (800 nm SP FESH0800, 900 nm LP dichroic DMLP900R, 900 nm LP FELH900, ThorLabs) coupled to a Princeton Instruments spectrometer (SCT 320) and liquid nitrogen cooled Princeton Instruments InGaAs detector (PyLoN-IR 1024/1.7). Fluorescence measurements were done with a beam-expanded 721 nm laser (10-500 mW, OptoEngine LLC) excitation light source and 800 – 1400 nm emission wavelength range. Solution-phase measurements were acquired in a 384 well-plate format (1 s exposure time, 1 mW laser power). Protein solutions (final concentration 40 μg/mL) or PBS control were incubated with (GT)_6_-SWCNTs (final concentration 5 μg/mL in 0.1 M PBS). For each time point, an aliquot of these incubation solutions was added to a well (40 μL total volume) and an initial fluorescence spectrum was acquired. 10 μL of dopamine was added to a final concentration of 200 μM prior to the second fluorescence acquisition. Fluorescence fold change was measured by taking the ratio of fluorescence intensities at 1200 nm between post- and pre-addition of dopamine spectra.

Similarly, surfactant-induced solvatochromism was performed by collecting nIR fluorescence spectra pre- and 1-minute post-addition of 0.5% (w/v) SDBS. Fluorescence fold change was defined as the ratio of integrated fluorescence intensity (800 to 1400 nm) between post- and pre-ddition of SDBS. Wavelength shift was measured relative to the wavelength of the (7,6) SWCNT chirality peak emission (initially 1130 nm) post-SDBS.

### 4.5 Kinetic model

Corona exchange kinetics were modeled by a system of simultaneous adsorption/desorption reactions. The model assumes that both ssDNA and protein adsorb/desorb reversibly to a fixed number of vacant SWCNT surface sites (Equations 1 and 2). Note that all modeling was done on a mass basis. This is in agreement with the general use of volume fractions in polymer thermodynamics.^52^ Here, we add the additional assumption that the biomolecules are of similar density. Modeling on a mass basis accounts for the widely varying molecular sizes between the two types of protein (HSA, 66.5 kDa, globular vs. FBG, 340 kDa, long) and ssDNA ((GT)_6_, 3.7 kDa). The time-dependent differential equations governing ssDNA desorption and protein adsorption are as follows:

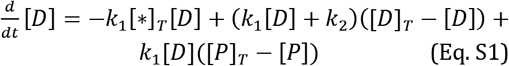

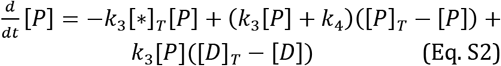

A least-squares regression was used to fit the model to fluorescence data and iterate model parameters. MATLAB 2019A’s ode15s solver was implemented to solve Equations S1 and S2 for free protein and DNA concentration curves given fit rate constants k_1_, k_2_, k_3_, k_4_, and the total concentration of open sites, [*]_T_. Relative error between the model fit and experimental data was calculated and averaged over all data points to yield mean relative error (MRE). Sensitivity analysis on initial conditions was performed to minimize this fit error. 48 unique combinations of rate parameter initial conditions were analyzed as inputs to the nonlinear least-squares solver (lsqcurvefit) in our MATLAB model. The optimal set of initial conditions for each protein was chosen as that which yielded a low MRE between fit and experimental data and a low standard error among fit parameters for each of the four protein concentrations. Each rate parameter was individually fit to each experiment, yielding 20 total fit parameters from each initial condition (Table S2). Final ssDNA and protein fit MRE were all < 5% (Table S2). Optimized initial conditions and resultant rate parameters for HSA and FBG are reported in Table S3.

Two alternative models were also attempted: in Model 2, protein was able to bind to surface-adsorbed ssDNA and in Model 3, protein was able to bind to surface-adsorbed protein. However, these models both produced significantly higher error in fits. Model 2 addressed the possibility of protein binding on top of ssDNA bound directly to the SWCNT. For FBG experiments fit with Model 2, most fits overestimated FBG adsorption and many fits displayed incorrect concavity for the ssDNA desorption. For HSA experiments fit with Model 2, the protein data was generally fit well but the ssDNA fits exhibited either a maximum or produced linear fits. Model 3 addressed the possibility of protein binding on top of protein bound directly to the SWCNT. For FBG experiments, Model 3 overestimated both protein adsorption and ssDNA desorption. For HSA experiments, Model 3 generally fit the protein data well, yet did not capture ssDNA dynamics as a function of concentration. Although the higher errors associated with Model 2 and 3 do not rule out these nanoscale mechanistic possibilities, the simple model of independent binding does overall fit the data much more closely between both protein and ssDNA curves within the same experiment, as well as binding dynamics as a function of varying concentration.

## ASSOCIATED CONTENT

### Supporting Information

Figures S1-S7 and Tables S1-S3 included with further experimental validation.

## ACKNOWLEDGMENT

We acknowledge support of an NIH NIDA CEBRA award # R21DA044010 (to M.P.L.), a Burroughs Wellcome Fund Career Award at the Scientific Interface (CASI) (to M.P.L.), the Simons Foundation (to M.P.L.), a Stanley Fahn PDF Junior Faculty Grant with Award # PF-JFA-1760 (to M.P.L.), a Beckman Foundation Young Investigator Award (to M.P.L.), and a DARPA Young Investigator Award (to M.P.L.). M.P.L. is a Chan Zuckerberg Biohub investigator. R.L.P., D.Y., and W.C. all acknowledge the support of NSF Graduate Research Fellowships.

## ABBREVIATIONS

ssDNA: single-stranded DNA
GT: guanine-thymine
SWCNT: single-walled carbon nanotube
HSA: human serum albumin
FBG: fibrinogen
PBS: phosphate-buffered saline
nIR: near-infrared
SDBS: sodium dodecylbenzenesulfonate

