## Supplemental Information for "Corona exchange dynamics on carbon nanotubes by multiplexed fluorescence monitoring" (from another version of this preprint: version 1, 2019-09-08)

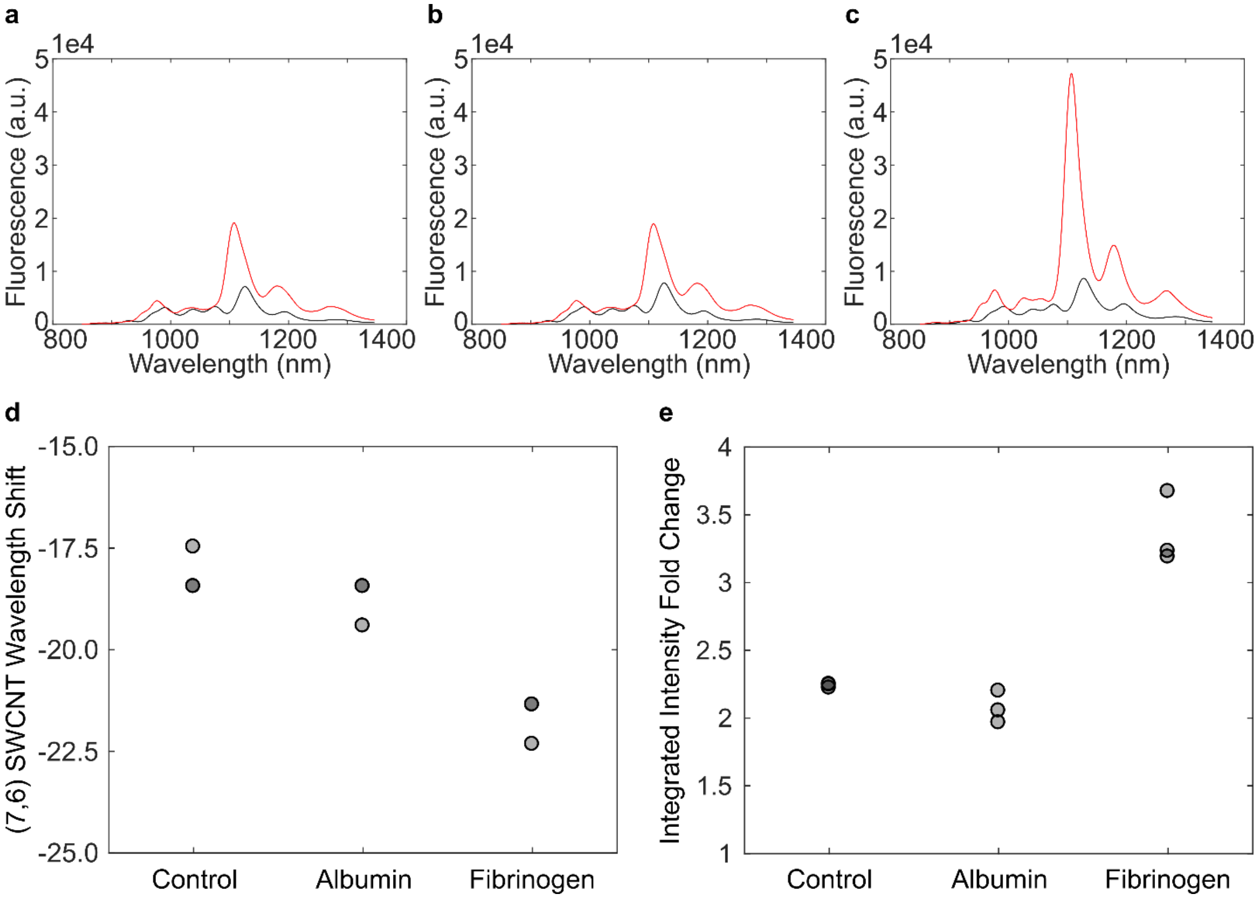


**Figure S1. Surfactant-induced fluorescence change of ssDNA-SWCNTs incubated with protein.** Near-infrared (nIR) spectra of 5 μg/mL $(GT)_6$-SWCNTs before (black) and after (red) addition of 0.5% (w/v) SDBS. **(b-c)** nIR spectra of 5 μg/mL $(GT)_6$-SWCNTs incubated with 40 μg/mL **(b)** albumin or **(c)** fibrinogen for 40 minutes before (black) and after (red) addition of 0.5% (w/v) SDBS. Change in **(d)** (7,6) SWCNT wavelength of peak emission and **(e)** integrated fluorescence intensity (800-1400 nm) observed 1 minute after addition of SDBS (N = 3). Nanosensor excitation was with 721 nm light.

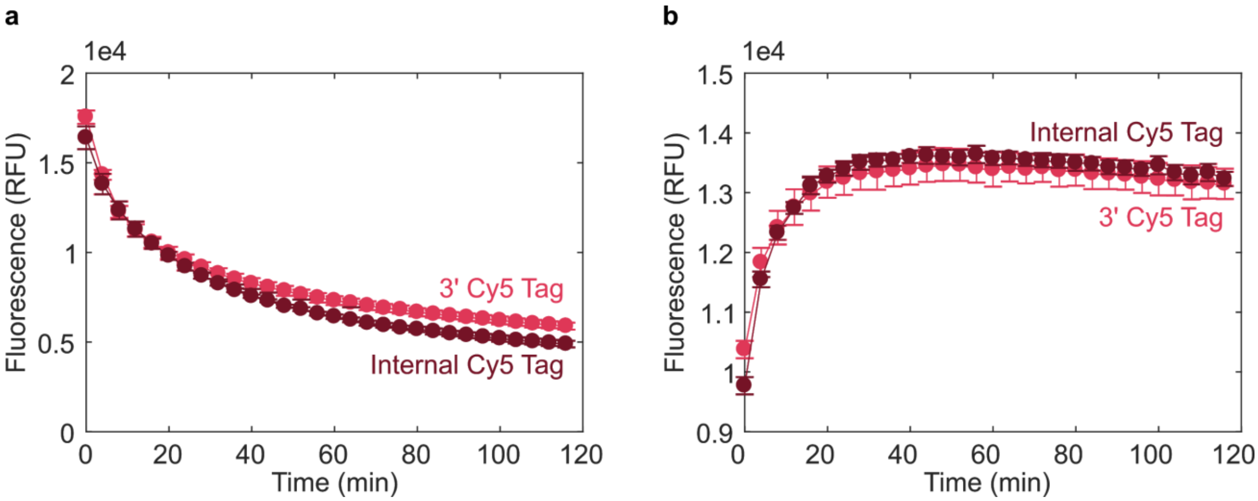


**Figure S2. Effect of Cy5 tag location on ssDNA adsorption and protein-induced desorption.** **(a)** Cy5 fluorescence tracking of 0.4 μM 3'- or internal-Cy5 tagged (GT)6 ssDNA added to 5 μg/mL unlabeled (GT)6-SWCNT. **(b)** Cy5 fluorescence tracking upon addition of 160 μg/mL fibrinogen, 3 hours post-incubation with (GT)6-Cy5. Error bars represent standard error between experimental replicates (N = 3).

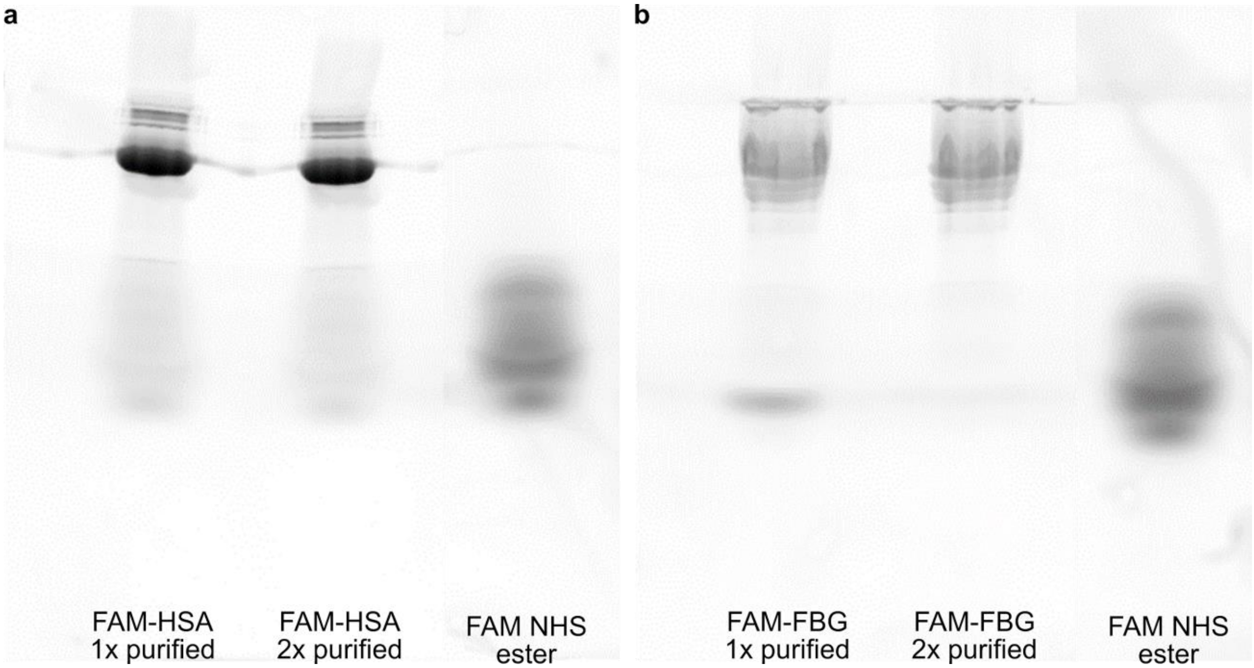


**Figure S3. Purification of labeled FAM-protein solutions.** 12% acrylamide (total monomer) gel images of FAM-labeled **(a)** albumin (FAM-HSA) and **(b)** fibrinogen (FAM-FBG) post-purification to assess amount of free FAM remaining.

**Table S1. Quantification of free FAM remaining in labeled FAM-protein solutions.** Molar percentages of FAM-protein and free FAM are calculated based on gel band intensity and protein degree of labeling for **(a)** FAM-HSA and **(b)** FAM-FBG.

**a**

| Sample | Band ID | Mole % |
|---|---|---|
| FAM-HSA, 1x pure | FAM-Protein | 69.7% |
| | FAM | 30.3% |
| FAM-HSA, 2x pure | FAM-Protein | 67.8% |
| | FAM | 32.2% |
| FAM | FAM | 100.0% |

**b**

| Sample | Band ID | Mole % |
|---|---|---|
| FAM-FBG, 1x pure | FAM-Protein | 70.5% |
| | FAM | 29.5% |
| FAM-FBG, 2x pure | FAM-Protein | 86.2% |
| | FAM | 13.8% |
| FAM | FAM | 100.0% |

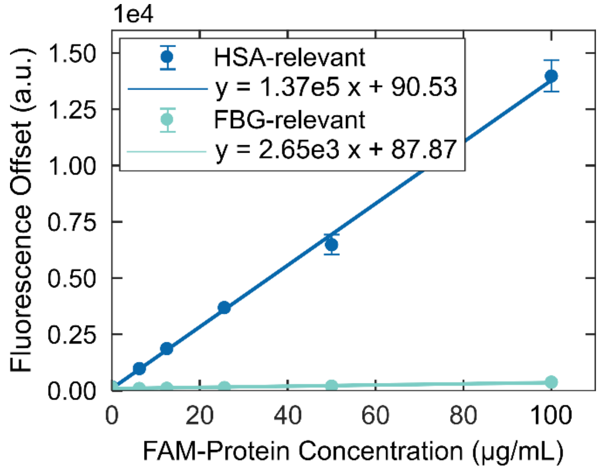


**Figure S4. Fluorescence offset due to free FAM binding to SWCNT surface.** FAM NHS ester was injected into solution with or without SWCNTs to quantify free fluorophore binding to the SWCNT. Free FAM concentrations tested were calculated based on molar percentage of free fluorophore at FAM-HSA or FAM-FBG concentrations between 0 and 100 μg/mL. Fluorescence offset was calculated as the difference between FAM fluorescence in the presence and absence SWCNTs. Offset is incorporated for all data conversion from fluorescence to concentration. Error bars represent standard deviation between technical replicates (N = 3).

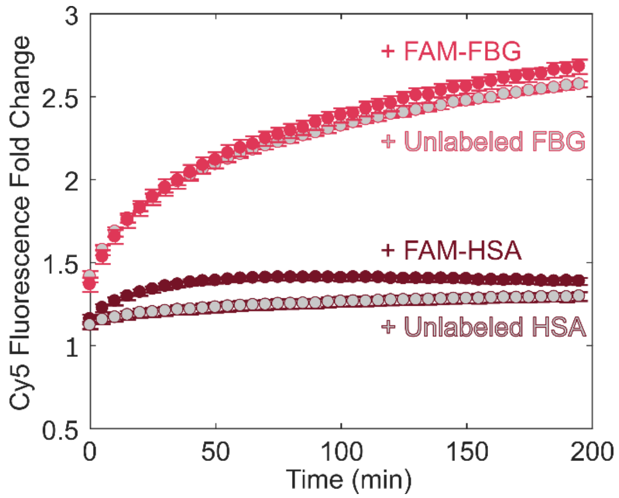


**Figure S5. Desorption of ssDNA-Cy5 from SWCNT induced by FAM-labeled vs. unlabeled protein.** Dequenching of (GT)6-Cy5 due to desorption from SWCNT upon addition of 40 μg/mL FAM-FBG or unlabeled FBG. Error bars represent standard error between experimental replicates (N = 3).

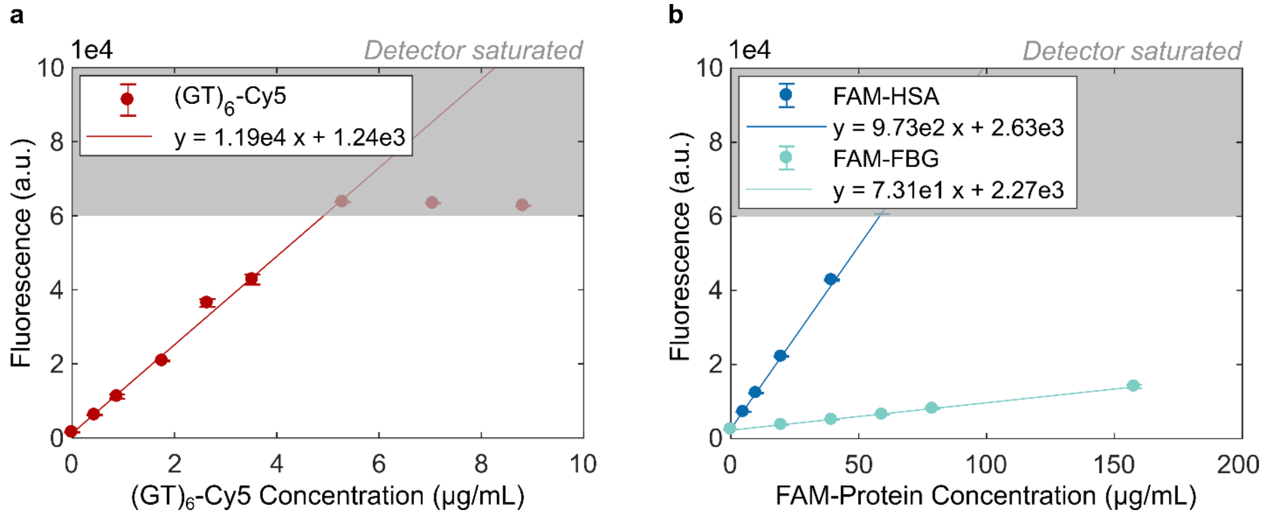


**Figure S6. Conversion of fluorescence to concentration.** Standard curves correlating fluorescence to fluorophore-labeled entity concentration for **(a)** Cy5-labeled $(GT)_6$ and **(b)** FAM-labeled proteins, FAM-HSA and FAM-FBG. Error bars represent standard deviation between technical replicates (N = 3).

**Table S2. Full kinetic model fit parameters and mean relative errors (MREs).**

| Protein | Concentration (μg/mL) | $k_1$ (mL μg$^{-1}$ s$^{-1}$) | $k_2$ (s$^{-1}$) | $k_3$ (mL μg$^{-1}$ s$^{-1}$) | $k_4$ (s$^{-1}$) | $[*]_T$ (μg mL$^{-1}$) | MRE protein | MRE ssDNA |
|---|---|---|---|---|---|---|---|---|
| Albumin | 5 | 1.54E-06 | 8.40E-06 | 9.44E-06 | 5.9E-03 | 364.60 | 1.21% | 0.25% |
| | 10 | 1.41E-06 | 1.18E-05 | 7.86E-06 | 8.1E-03 | 525.60 | 0.83% | 0.90% |
| | 40 | 1.10E-06 | 1.34E-05 | 8.87E-06 | 1.1E-02 | 514.26 | 0.35% | 3.01% |
| | 60 | 1.22E-06 | 2.07E-05 | 9.34E-06 | 1.2E-02 | 483.76 | 0.41% | 2.53% |
| Fibrinogen | 40 | 1.15E-06 | 4.27E-05 | 1.38E-05 | 2.6E-03 | 619.82 | 4.71% | 2.96% |
| | 60 | 2.18E-06 | 6.64E-05 | 1.69E-05 | 3.7E-03 | 485.72 | 2.98% | 3.47% |
| | 80 | 2.09E-06 | 7.65E-05 | 1.48E-05 | 5.4E-03 | 562.90 | 1.37% | 4.14% |
| | 160 | 2.30E-06 | 9.09E-05 | 1.18E-05 | 9.2E-03 | 617.74 | 0.74% | 4.74% |

**Table S3. Sensitivity analysis in kinetic fitting and resultant optimized initial conditions.**

| | $k_1$ (mL μg$^{-1}$ s$^{-1}$) | $k_2$ (s$^{-1}$) | $k_3$ (mL μg$^{-1}$ s$^{-1}$) | $k_4$ (s$^{-1}$) | $[*]_T$ (μg mL$^{-1}$) |
|---|---|---|---|---|---|
| **Range of Initial Conditions Tested** | 1.0E-03 | 1.0E-02 | 1.0E-03 | 1.0 | 1000 |
| | 1.0E-04 | 1.0E-03 | 1.0E-04 | 0.1 | 100 |
| | 1.0E-05 | | | | |
| **Albumin Optimized Initial Conditions** | 1.0E-04 | 1.0E-02 | 1.0E-03 | 1.0 | 100 |
| **Fibrinogen Optimized Initial Conditions** | 1.0E-04 | 1.0E-02 | 1.0E-04 | 0.1 | 100 |

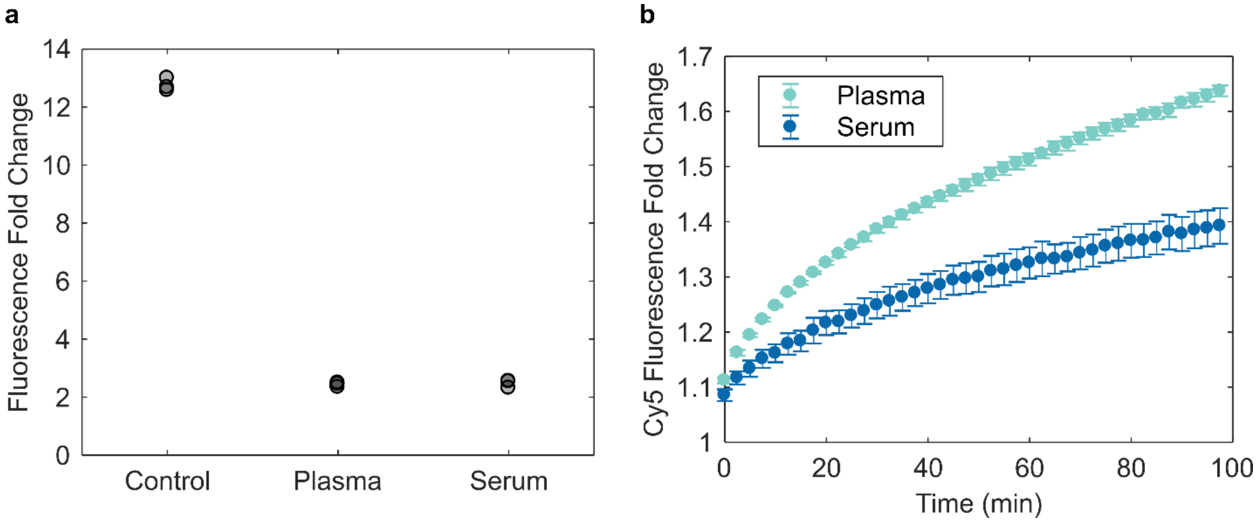


**Figure S7. Comparison of plasma- vs. serum-induced effects on ssDNA-SWCNTs. (a)** Change in 1200 nm fluorescence intensity of 5 μg/mL $(GT)_6$-SWCNTs, pre-incubated for 40 minutes with PBS, plasma, or serum normalized to 40 μg/mL total protein concentration, before and after addition of 200 μM dopamine (N = 3). Nanosensor excitation was with 721 nm light. **(b)** $(GT)_6$-Cy5 fluorescence fold change upon desorption from 5 μg/mL SWCNTs induced by addition of plasma or serum, 40 μg/mL total protein concentration. Error bars represent standard error between experimental replicates (N = 3).
